# Natural weed seed predators reduce crop yield loss due to weeds by 20% in cereal fields

**DOI:** 10.1101/2024.09.05.611395

**Authors:** Lucile Muneret, Benjamin Carbonne, Bruno Chauvel, Alexandre Dosset, Chantal Ducourtieux, Nicolas Henon, Emeline Felten, Emilien Laurent, Annick Matejicek, Sandrine Petit

## Abstract

While many herbicide active substances have been banned and weed infestation is a major threat to crop productivity, it is still unknown to what extent natural weed control provided by seed predators can help farmers manage weed communities. We aim to quantify the contribution of seed predators to crop productivity through weed control and to evaluate whether the magnitude of their influence depends on farming systems such as conservation agriculture. We set up 112 seed predator-exclusion cages in 28 cereal fields in France (14 pairs of conservation and conventional agriculture fields), surveyed weed emergence and biomass, measured crop yield and sampled the main seed predators: carabid beetles and rodents. We found that seed predators’ activity reduces the yield loss due to weeds by 20%. By extrapolation, it represents an economic gain of 285€/ha. However, the yield loss remains at 60% below the maximum crop yield potential reached in the absence of weeds. Moreover, conservation agriculture enhances weed control, but this does not translate into increased crop yield. This study demonstrates the tangible importance of considering seed predators for weed control but highlights the need to combine this approach with weed control practices or to substantially redesign cropping systems to enhance the beneficial effects of biodiversity on crop productivity.

**Significance statement:** Although weed biomass is the main driver of decreasing the crop productivity worldwide and the use of herbicides is massively disparaged, we have not yet quantified the role played by on-farm biodiversity to control weeds. Using an experimental design set up in 28 commercial cereal fields in France, we showed that weed seed predators reduce crop yield loss due to weeds by 20%. By extrapolation, it represents an economic gain of 285€/ha. However, in the absence of any other weed control practices, the yield loss remains at 60% below the maximum crop yield potential reached in the absence of weeds. This study demonstrates the quantitative importance of considering seed predators to design pesticide-free systems.

## Introduction

Crop yield highly depends on synthetic fertilizers and pesticides in industrialized countries (Tilman, 2002). However, due to their negative impacts on the environment and biodiversity worldwide, their use is increasingly questioned and many pesticides are regularly prohibited (Jacquet et al., 2022; Chauvel et al., 2022). As an alternative paradigm, ecological intensification of agriculture means that ecological processes supporting intermediate ecosystem services can replace —at least partially— the use of external inputs to enhance crop yield (Bommarco et al., 2013, Dainese et al., 2019). Yet, this new paradigm comes with its share of uncertainties. First, agrobiont biodiversity has suffered from agricultural intensification for decades altering the provision of efficient regulating ecosystem services such as pollination or pest control potential (Rigal et al., 2023; Sanchez-Bayo et al., 2019). Second, aiming to enhance biodiversity in agricultural fields via agroecological practices or maintaining seminatural habitats could also result in more damaging crop pests or weeds detrimental to crop yield (Muneret et al., 2018; Delaune et al., 2021). Therefore, harnessing biodiversity to provide efficient regulating services that limit pest outbreaks and foster crop yield through the implementation of biodiversity-friendly practices is still a huge challenge.

Despite herbicide use, weeds are the main biotic threat to crop yield (Oerke et al., 2006). The biomass of dominant weeds is a strong driver of yield loss in arable fields (Adeux et al., 2019). It might be especially significant under ecological intensification context because of the weed seedbank persistence and its species diversity which can respond to different environmental conditions. Many macro-organisms like carabids, ants, voles and granivorous birds have been identified as weed seed predators (Sarabi, 2019). They can reduce weed emergence by 38% and weed biomass by 81% in fallow plots (Blubaugh & Kaplan, 2016). Due to compensation mechanisms of weeds leading to better growth and higher seed production when there are fewer weeds, it is still unknown whether weed seed predators really impact crop productivity through weed control (Pannwitt et al., 2021). Such knowledge is lacking because, instead of mentioning a vague effect of biodiversity on weed dynamics, we could quantify what we can really expect about natural weed control and be clear about its concrete effects with regards to farmers.

The implementation of biodiversity-friendly practices or systems is expected to enhance biodiversity and regulating ecosystem services (Wittwer et al., 2021). However, implementing an agroecological system can also lead to crop yield loss because regulating services might take time to become effective or not offset the absence of external inputs to reach productivity goals. For example, conservation agriculture is generally beneficial for biodiversity but the increased complexity of trophic interactions can dilute their effect on the provision of pest control services (Henneron et al., 2015; Carbonne et al., 2023). Moreover, the conservation agriculture is on average 5% less productive than conventional agriculture (Pittelkow et al., 2015). Measuring the efficiency of alternative systems such as conservation agriculture in enhancing regulating services is still necessary to guide the agroecological transition.

Here, we conducted an in-field experiment establishing the contribution of seed predators to crop yield. First, we hypothesized that seed predators’ activity (invertebrates and vertebrates) reduces weed emergence and biomass, which in turn increases crop productivity. Second, we assumed that conservation agriculture increases invertebrate and vertebrate seed predators, strengthening the positive effect of weed control on crop yield.

We set up four types of seed predators-exclusion cages in 28 commercial arable fields in France (14 pairs composed of one conservation and one conventional field, Figure 1). In the first cage, only wheat was sown, i.e. without weeds and giving no access to seed predators, it represents the maximum crop productivity potential (hereafter, it is considered as the cage “*Control”*). In the second cage, wheat and weeds were sown, without any access to seed predators, it represented the weed harmfulness by preventing all predators from accessing to the weed seeds (hereafter the cage “*No Predation”*). The third cage included wheat, weeds and gave access to invertebrate seed predators only (named as “*Partial Predation*”) and the fourth cage included wheat, weeds and gave access to invertebrates and vertebrates seed predators (hereafter “*Total Predation*”). The soil contained in the cages was previously sterilized before setting. In spring 2021, 234 weed seeds from nine species were sown to mimic natural seed rain and then 20 wheat seeds were sown in November, the usual sowing period for farmers in the study area.

**Figure 1.**
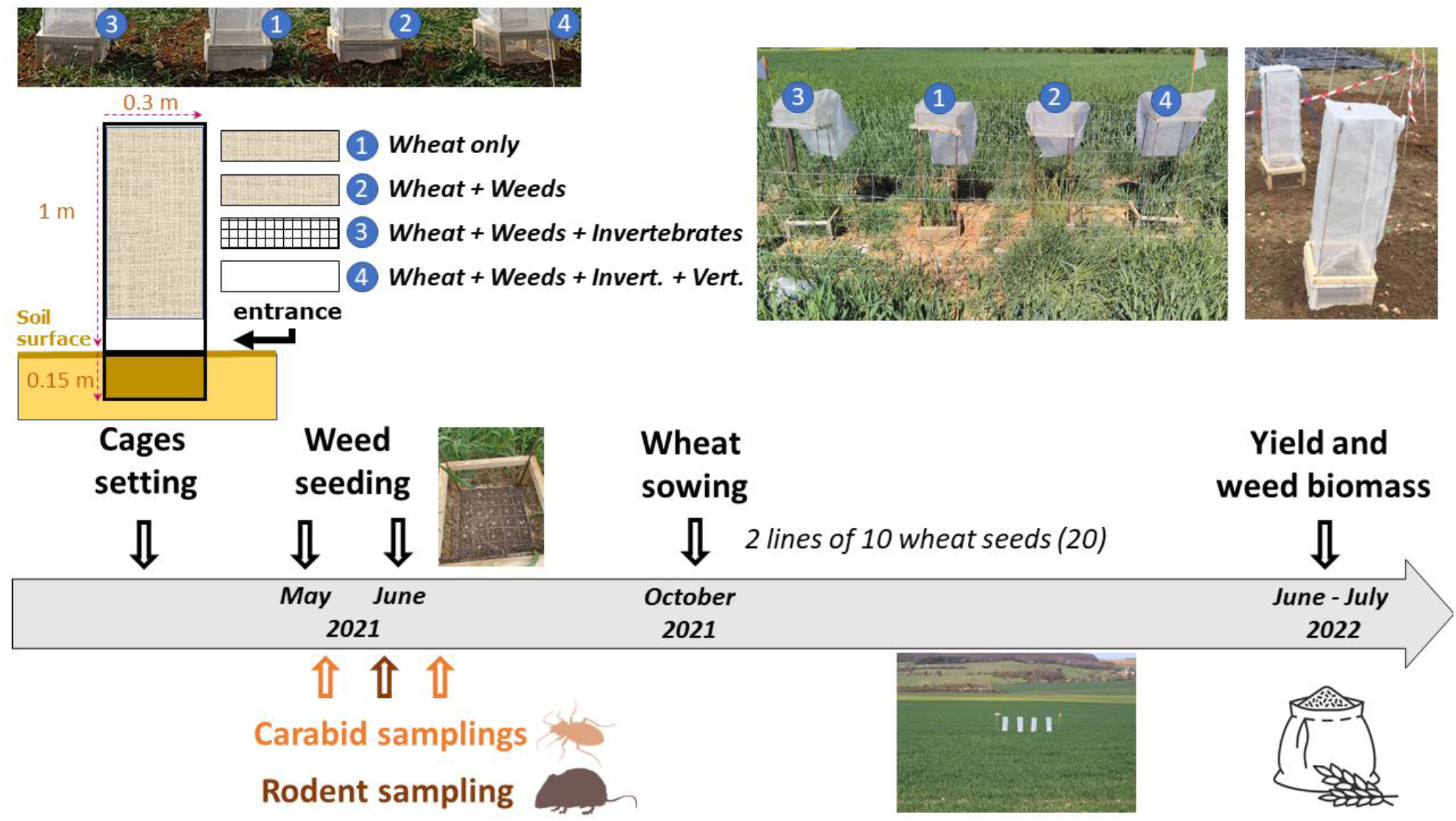
Overview of the timeline of the experiment, setting and description of the four modalities. (1) Yield Max Potential, (2) No Predation, (3) Partial Predation and (4) Total Predation.

We surveyed both wheat and weed growth until harvest in July 2022. In parallel, we sampled invertebrates and vertebrates (especially carabid beetles and rodents) in spring-summer 2021when they can eat the seeds following seed rain.

## Results

Our experiment demonstrates that wheat productivity was reduced on average by 80 % in the presence of weed seeds if no seed predators could access the seeds, but that this yield loss value was on average 60% when invertebrate and vertebrate seed predators had access to weed seeds. We were able to demonstrate the links underpinning this result, with a significant reduction of weed biomass in the presence of seed predators, and a significant impact of weed biomass at harvest on wheat productivity.

### Responses of crop productivity and weed biomass to weed seed predators’ exclusions

The crop yield measured in all the four seed predators-exclusion cages varied significantly except between the two cages *Partial Predation* and *Total Predation* (paired t-test based on a total of 112 observations, Figure 2A; Supplementary Table 1). The maximum crop yield was reached in the cage *Control*, with 73.3g of wheat grains on average (Standard Deviation (SD) 59.2). In contrast, the cage *No Predation* produced 14.7g (SD 24.5) of wheat grains meaning a reduction of 80% of the crop productivity relatively to the cage *Control*. Cages *Partial Predation* and *Total Predation* produced respectively 39.8g (SD 40.8) and 29.6g (SD 35.6) of wheat grains on average. Crop productivity was therefore reduced by 46% and 60% respectively in comparison to the cage *Control*. This also showed a slight negative influence of vertebrate predators on crop productivity. Alternatively, crop productivity is multiplied by 2.7 and 2.0 in the cages *Partial Predation* and *Total predation* respectively with regard to the cage *No Predation*.

**Figure 2.**
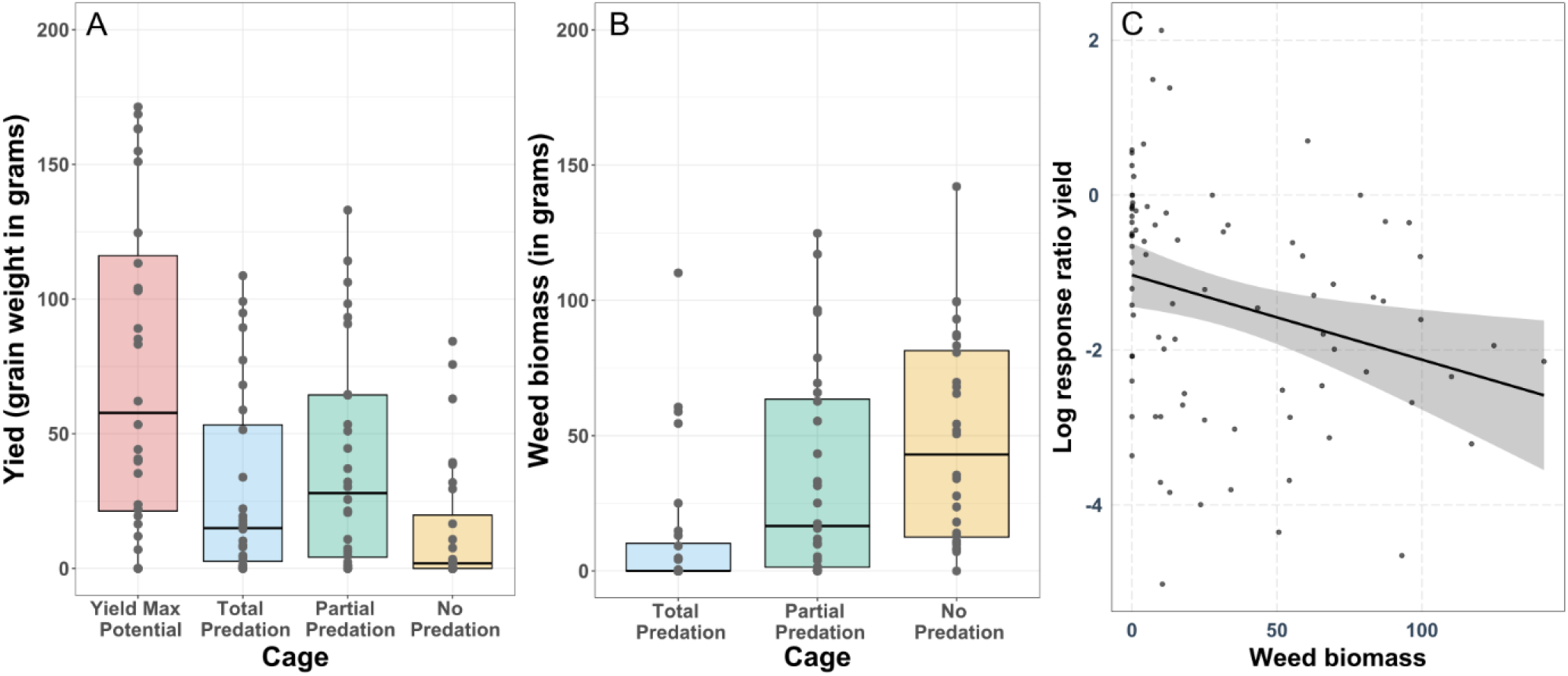
Responses of (A) crop productivity and (B) weed biomass to weed seed predators’ exclusions and (C) relationships between weed biomass (in grams) and response ratio of the crop productivity (a low log response ratio indicates an important yield loss).

By extrapolation, at the hectare scale and without any other weed control measures than natural weed seed predation, crop productivity reached 8.1t/ha, 1.6t/ha, 4.4t/ha and 3.3t/ha in the cages *Control, No predation, Partial predation* and *Total Predation* respectively. Based on these experimental conditions, considering the price of winter soft wheat over the 10 last years (July 2014 to July 2023, i.e. wheat price at 167€/t, FranceAgriMer), seed predator gains reach 285€.hectare^-1^.-year^-1^ (price difference per hectare between the cage *No Predation* and the cage *Total Predation*).

In the three cages that contained weed seeds (84 observations, excluding the cages *Control)*, the paired t-test revealed significant differences of weed biomass at harvest (Figure 2B; Supplementary Table 1). The cage *Total Predation* had a significantly lower level of weed biomass than the two other ones i.e. *Partial Predation* and *No Predation*. The cage *Partial Predation* had an intermediate level of weed biomass and the cage *No Predation* had the highest level of weed biomass but these two latter were not significantly different.

### Relationship between weed biomass and crop yield loss

To evaluate the effect of weed biomass on crop yield loss, we performed a linear regression (N=84) between weed biomass and the crop yield loss. The indicator describing the crop yield loss used was the response ratio of the crop productivity between each cage and the cage *Control*: (i.e. ln ((Yfk +1) / (Yfc+1)) with *Y* as the crop productivity, *f* as a *field, k* as a cage either *No Predation* or *Partial Predation* or *Total Predation* and *c* as the cage *Control*; Figure 2C). We detected a slight negative relationship between weed biomass and the response ratio of crop productivity (estimate = -0.01, t (82) = -2.614, p-value = 0.01) meaning that when the weed biomass is low, crop productivity is closer to the maximum crop productivity potential measured in the cage *Control*.

### Effect of conservation agriculture and seed predators on weed biomass and crop yield loss

Two Piecewise Structural Equation Models (“PSEM”) revealed the links between farming systems, seed predators, weed emergence, weed biomass and crop yield loss for the cages *Total Predation* and *Partial Predation* independently (each one based on 28 observations; Figure 3, conceptual framework). When both carabids and rodents were considered (cages *Total Predation*) the PSEM performed poorly at explaining the crop yield loss (R^2^ = 0.04; Supplementary Table 3). All the intermediate links between farming systems, predator biomass (carabid beetles and rodents), weed emergence, weed biomass and crop productivity response ratio were consistent with the hypotheses but insignificant (Figure 3). Specifically, conservation agriculture tended to increase carabid and rodent biomasses, that tended to reduce weed emergence. Weed emergence tended to increase weed biomass that in turn reduced the crop yield loss. When only carabids were considered (cages *Partial Predation*) causal links explained the crop yield loss better than in the previous model (R^2^=0.23; Figure 3, Supplementary Table 3). All the links detected were in accordance with hypotheses. Conservation agriculture tended to increase carabid biomass (ns, standardized coefficient (std. est.) = 0.29, R^2^=0.09), carabid biomass reduced weed emergence (std. est. = -0.52, R^2^=0.27) and weed biomass reduced crop yield loss (std. est. = -0.48, R^2^=0.23) in *Partial Predation* cages. We also detected a significant direct negative link between conservation agriculture and weed biomass (std. est. = -0.49, R^2^=0.22) leading to a positive cascading effect of conservation agriculture on the crop yield through the weed biomass reduction (Supplementary table 4).

**Figure 3.**
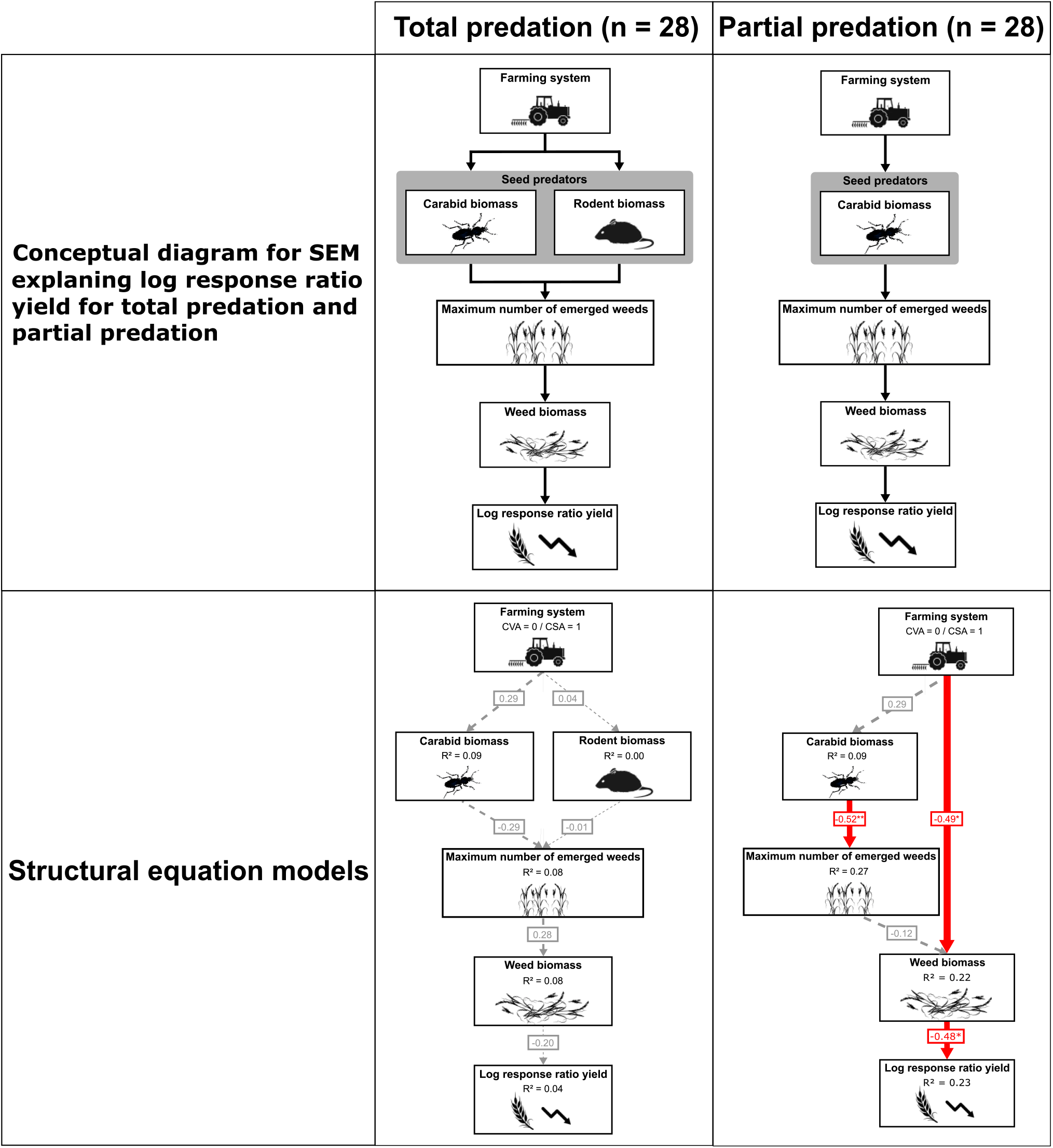
Conceptual frameworks and results of both PSEM used to investigate how farming systems affect crop productivity through natural enemies and weed development. A low log response ratio indicates an important yield loss. Solid red arrows represent significant negative links and dashed grey represent insignificant links. Standardised coefficients are shown for each path. The numbers of asterisks indicate the level of significance (p<0.1, *p<0.05, **p<0.01,***p< 0.001).

## Discussion

We quantified for the first time the effect of seed predators on crop productivity, mediated by a reduction of weed biomass. We found that crop yield loss due to weeds is reduced by 20% when all the seed predators have access to and can consume seeds. This regulation by seed predators would be equivalent to an economical gain of 285€/hectare-year^-1^, under the assumption that our cage-scale results can be extrapolated to the field scale. However, under this condition, crop yield loss due to weeds still amounted to 60% of the maximum crop yield potential. Moreover, conservation agriculture tended to increase weed control by enhancing seed predator’s biomass, which reduced weed emergence but did not translate into higher crop yield (Supplementary Figure 1, Supplementary Table 5). This work shows the importance of considering weed seed predation for field management, but also the necessity to either combine it with other technics or drastically redesign agricultural management to boost the seed predator’ activity (considering potential negative effect of vertebrate seed predators on crop productivity).

In accordance with our expectations, crop productivity responded positively, and weed biomass negatively, to the open access of weed seed predators in the cages (Figure 2). The reduction of crop yield loss due to weeds by 20% and allowed by seed predators means that weed compensation mechanisms, i.e. isolated weed plants can provide more biomass and then more seeds than an abundant weed community, do not balance the effect of seed predators on the limitation of weed biomass. This is the first demonstration of the concrete effect of seed predators on crop yield. Another work already showed that their effect on the reduction of weed emergence could reach 38% in fallow plots (Blubaugh et Kaplan, 2016). However, when invertebrate and vertebrate seed predators contribute to weed seed removal, crop productivity remains at 60% below the crop productivity potential. This aligns with results from modelling approaches showing that seed predators need to consume at least 78% of the seeds at peak seed shedding to suppress *Alopecurus myosuroides* growth (Daouti et al., 2022). This threshold was clearly not reached here and shows the necessity of combining multiple measures to control weeds efficiently.

We carried out a rough economic evaluation of the service provided by seed predators, they saved 285€/hectare-year^-1^ in terms of crop yield. To our knowledge, this is the first attempt of valuating weed control provided by seed predators. Previous studies showed that pest control provided by bats and birds can range between 44$ and 730$/hectare-year^-1^ according to study, location and agroecosystems (Kellermann et al., 2008 ; Maas et al., 2013 ; Karp et al., 2013). Our estimation is in this range. The plus of this effort is to reconsider the way of measuring productivity in fields including the contribution of biodiversity (Seppelt et al., 2020). However, this economic valuation has to be taken with caution because of price fluctuations and because maximizing the provision of regulating services does not systematically lead to meet biodiversity conservation goals (Adams et al., 2014; Kleijn et al., 2015).

The comparison of cages giving access to either invertebrate predators only or invertebrate and vertebrate predators did not turn into a significant difference of crop productivity. However, the yield loss was lower when only invertebrate predators are involved (−46%) than when invertebrate and vertebrate predators are at play (−60%). This corroborates that vertebrates, notably voles, are seed predators but also have negative effects on crop plants (Krijger et al., 2017). Considering a way of controlling them in addition to weed control is necessary to limit the crop yield loss.

Conservation agriculture had a positive but insignificant effect on seed predators’ biomass such as carabid beetles and rodents. First, the result is due to the choice of dataset used in the PSEM, we considered granivorous and omnivorous species simultaneously while a previous work on this dataset found a positive effect of conservation agriculture on the activity density of granivorous carabid species, as well as on other taxonomic groups (Carbonne et al., 2023). Moreover, we detected a clear effect of conservation agriculture on the reduction of weed biomass. This link suggests that other taxonomic groups not included in the PSEM can contribute actively to weed control (Sarabi, 2019). Implementing conservation agriculture stimulates weed control services but not at level preventing yield loss due to weed biomass interference.

## Material and methods

### Experimental design

Our experiment was carried out in 14 pairs of cereal fields in a cropland-dominated area around Dijon in East of France (47°19′19.369″ N, 5°2′29.328″ E). Pairs were composed of one field conducted under conservation agriculture and the second referred as conventional agriculture. They covered a gradient of proportional cover of conservation agriculture in a buffer of 1 km radius around fields (a range from 3.1% to 37.5%). This metric was not correlated with the proportion of semi-natural habitats. The landscape was documented using the French Graphic Parcel Register, superimposed on the CORINE Land Cover map 2000 in ArcMap 10.8. The land use and the farming system of the parcels were also recorded through field and farmer surveys. The mean distance between two nearest sampled field pairs was 9.6 km. All field sizes ranged from 1.94 to 38.7 ha, with the exception of two smaller experimental fields from an experimental site. In each field, we set up four cages with different weed seed predator-exclusion treatments at 20–50 m from the field margins except for the two experimental fields (i.e. 112 cages in total). Cages were aligned, spaced 30 cm apart and all together enclosed by a wire fence positioned about 1m around the line of cages to protect from wild animals’ damages. No agricultural intervention was done within the fence and farmers applied their own technical routes in the field.

The cages were hand-made and composed of one buried and one aerial part. The buried part was a box measuring 0.30 × 0.30 × 0.15 m (L×W×H). The four edges were made of wood and the bottom was made of a perforated steel sheet (diameter 2 mm) to make water exchange possible with the field soil. The boxes were filled with sterilized sandy loam soil (®SONOFEP company). The boxes were buried into the soil to have their soil surface at the same level than the soil surface of the field (Figure 1). The aerial part was framed by a structure made of 8 mm steel bars having a height of 1.0 m. The difference between the four cages is in the 0.15 m just above the soil. This part was bordered by wood frames that were either closed by a plastic net giving no access to seed predators to the cage (the cages *Control* and *No predation* were like this), or by a chicken wire giving (diameter 0.012 m) only access to invertebrate predators or opened giving access to both vertebrate and invertebrate predators. Above this 0.15 m part, all the cages were covered by a plastic net (diameter 600 µm) to have the same microclimatic conditions in the cages independently of the treatment.

There were four cages. First, the cage with wheat, excluding weeds and seed predators, measures the maximum crop productivity potential, it is referred as the cage *Control*. Second, the cage with wheat and weeds and excluding totally seed predators, measures weed harmfulness to crop productivity, it is the cage *No Predation*. Third, the cage with wheat and weeds and giving access to invertebrates measures the control of weed harmfulness by invertebrates, it is the cage *Partial Predation*. Fourth, the cage with wheat, weeds and giving a total access to predators measures the control of weed harmfulness by invertebrates and vertebrates, it is the cage *Total Predation*.

Finally, the cages were set up in late March early April 2021 to have the minimum impact on the field crop. All the chronology of the experiment is reported in Figure 1 (settings and all the sampling dates were reported in the supplementary table 6). In the three cages including weeds, we placed 104 seeds from four annual weed species (i.e. *Geranium dissectum*; *Stellaria media*; *Veronica hederifolia*; *Viola arvensis* – N=26 for each species) evenly distributed in a grid at the soil surface of the cage in Mid-May. In Mid-June, we also evenly distributed 130 seeds from five annual weed taxa (i.e. *Alopecurus myosuroides*; *Galium aparine*; *Anisantha sterilis*; *Lolium* sp.; *Cyanus segetum* – N=26 for each species). Weed species were chosen because either they are defined as harmful by farmers in the area or because they are a well-studied biological model for weed control (*V. arvensis*). The seeds were placed into the cages when their natural seeds’ rain occur naturally in the area. In the four cages, 20 wheat seeds were sown along two lines spaced 0.17 m apart (as in the field on average) in the middle of the cage at a depth of 0.02 m manually in November. It corresponds to a density of 222 wheat seeds/m^2^ what is a probable density applied by farmers in France.

### Plant surveys

Weeds were surveyed nine times throughout the cropping season to remove each unsown plant. We counted the number of emerged plants. At harvest in late-June 2022, we cut by hand all the vegetation of the cage just above the soil surface and measured the dry biomass of the weeds and the wheat. We also measured the number of wheat grains as well as their weights to assess the crop yield of each cage. Plant biomass was obtained by drying in an oven at 80°C during 72 hours.

### Weed seed predator samplings

Carabids (Coleoptera: Carabidae) were sampled using three pitfall traps per field. Each trap was composed of a plastic beaker (10 cm height, 8 cm diameter), filled with 150 mL of a preservative solution of salt-saturated water and a few drops of odourless soap. A plastic cover was placed above each trap to avoid rain flooding. The traps were exposed for 7 days twice in May and June 2021 (Sampling dates in the supplementary table 6). Carabids were identified at the species level (Coulon et al., 2011). Carabids were assigned to a trophic group: carnivorous, omnivorous or granivorous (Homburg et al., 2013) and we only kept omnivorous and granivorous for further analyses. The total number of omnivorous and granivorous carabids collected in each field corresponds to the measure of activity density (abundances of both sampling sessions were summed). The number of specimens per species was multiplied by its specific body mass (*Body mass = 0*.*0237 × Body size*^*2*.*7054*^ ; Barnes et al. 2014). The body size of each carabid species was extracted from the literature (Appendix S1).

Rodents were captured using ‘INRA traps’ (16 cm long, 5 cm height, 5 cm width — BTTM company) connected to 500 mL boxes containing hay, a gel water ball and a food pellet (beef-based cat food and peanut butter) to meet the needs of the captured individuals. The traps were placed in two lines of 25 traps spaced at 3 m intervals. The two lines were placed near a tractor wheel path and separate by 20–30 m. The traps were exposed for three consecutive days and the presence of captures was checked every 24 h. Rodent sampling on all fields was conducted over a single period between late May and early June. Captured individuals were carefully identified, weighed, marked and released directly. All captures for the 3 days were pooled to obtain a total abundance and multiplied by the average individual body mass measured in field, this provides the rodent biomass.

### Statistical analyses

First, to compare the crop yield (absolute values) and weed biomass between the types of cages, we used paired t-tests (112 observations and 84 observations, respectively). Second, to calculate economic gains of natural weed control due to increased wheat productivity at the hectare scale, we multiplied the crop yield obtained in the cage by 11111.1 because the area in cages was 0.09m^2^ and because wheat seed were sowed at a commercial density to mimic growers’ sowing. Tons of wheat grains obtained were then multiplied by the mean price of a ton of the wheat grain between July 2014 to July 2023 (FranceAgriMer). Third, to examine the effect of weed biomass on crop yield loss, we ran a linear model including crop yield loss as the response variable, weed biomass as the predictor. Crop yield loss used corresponded to the response ratio of the crop yield between each cage and the cage *Control*: (i.e. ln ((Yfk +1) / (Yfc+1)) with *Y* as the crop productivity, *f* as a *field, k* as a cage either *No Predation* or *Partial Predation* or *Total Predation* and *c* as the cage *Control*). Fourth, two PSEM were developed to investigate causal links between farming system (conventional vs. conservation agriculture), seed predator biomass, weed emergence, weed biomass at harvest and crop yield. One PSEM was based on data from cages with *Partial Predation* and invertebrate seed predator communities. The second one was based on data from cages with *No Predation* and vertebrate and invertebrate communities. Homoscedasticity and normality of all model residuals were visually checked. The overall fit of both PSEM were assessed with D-separation tests n Fisher’s C statistic (Lefcheck, 2016) and AIC. All the analyses were performed using R-Core team (V 4.3.1) and PSEM were performed by the package piecewiseSEM (Lefchek, 2016) and direct and indirect links summarized using semEff (Murphy, 2023).

## Acknowledgements

This research was funded by the EIP-AGRI project RegGAE (funds from the EU Rural development 2014-2020 for Operational groups and the Region Bourgogne-Franche-Comté). We thank very much all the farmers who let gave us access to their fields and agreed to let us carry out our experiment. We also thank Stéphane Cordeau, Guillaume Adeux and all the colleagues of the Pole “Gestion durable des Adventices” for all the advices they provided when we conceived the study. Finally, we thank Rodolphe Hugard, Mathieu Siol, Charly Pouillot, Luc Biju-Duval, Anne-Lise Goumon, Philippe Aubert and Pascal Farcy who help us to build up the experimental design, the cages and contributed to the surveys.

## Data availability

Data are available on data.gouv.fr: Muneret, Lucile; Carbonne, Benjamin; Matejicek, Annick; Laurent, Emilien; Felten, Emeline; Henon, Nicolas; Ducourtieux, Chantal; Dosset, Alexandre; Chauvel, Bruno; Petit, Sandrine, 2024, “QUANTIF - Crop yield loss due to weed biomass, seed predators and farming systems”, https://doi.org/10.57745/2ZTMGR, Recherche Data Gouv, V1. Scripts are available at git@github.com:l-muneret/QUANTIF.git.

## Authors’ contributions

LM and SP conceived the study. All the authors contributed to build the experimental design, collected and managed the data. LM and AD analysed the data. AD wrote the open access scripts. LM wrote the first draft and all the authors contributed significantly to the final version of the manuscript.

## Supplementary materials

## Appendix 1

**References used to determine carabid identity.** *References used for assessing the body length of the carabid community: the mean value between the minimum and maximum values.

---

**Supplementary table 1.** Paired t-tests for comparison of crop yield and weed biomass between the different types of cages.

| Type | Comparison | t statistic | p value | Confidence intervals | df |
| --- | --- | --- | --- | --- | --- |
| Yield | Partial Predation - Total Predation | 1.34 | 0.19 | (-5.40 ; 25.83) | 27 |
| Yield | No predation - Total Predation | -2.95 | 0.01 | (-25.28 ; -4.56) | 27 |
| Yield | Control (Max Crop yield) - Total Predation | 4.92 | 0.00 | (25.45 ; 61.79) | 27 |
| Yield | Partial Predation - No predation | 4.17 | 0.00 | (12.77 ; 37.50) | 27 |
| Yield | Partial Predation - Control (Max Crop yield) | -4.40 | 0.00 | (-48.98 ; -17.82) | 27 |
| Yield | Control (Max Crop yield) - No predation | 6.45 | 0.00 | (39.93; 77.14) | 27 |
| Weed biomass | Partial Predation - Total Predation | 1.34 | 0.19 | (-5.40 ; 25.83) | 27 |
| Weed biomass | No predation - Total Predation | -2.95 | 0.01 | (-25.28 ; -4.56) | 27 |
| Weed biomass | Partial Predation - No predation | 4.17 | 0.00 | (12.77; 37.50) | 27 |

**Supplementary table 3.**
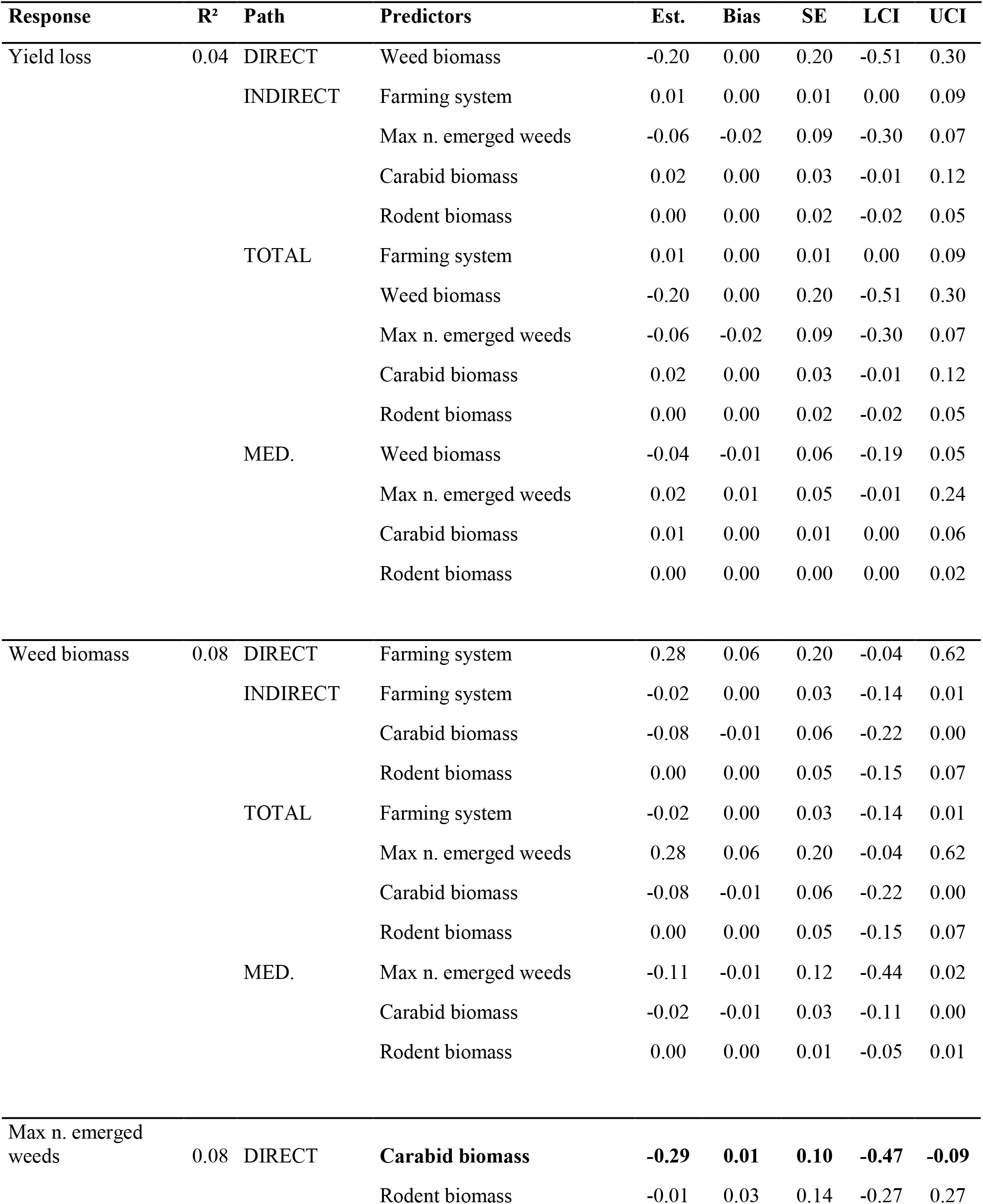
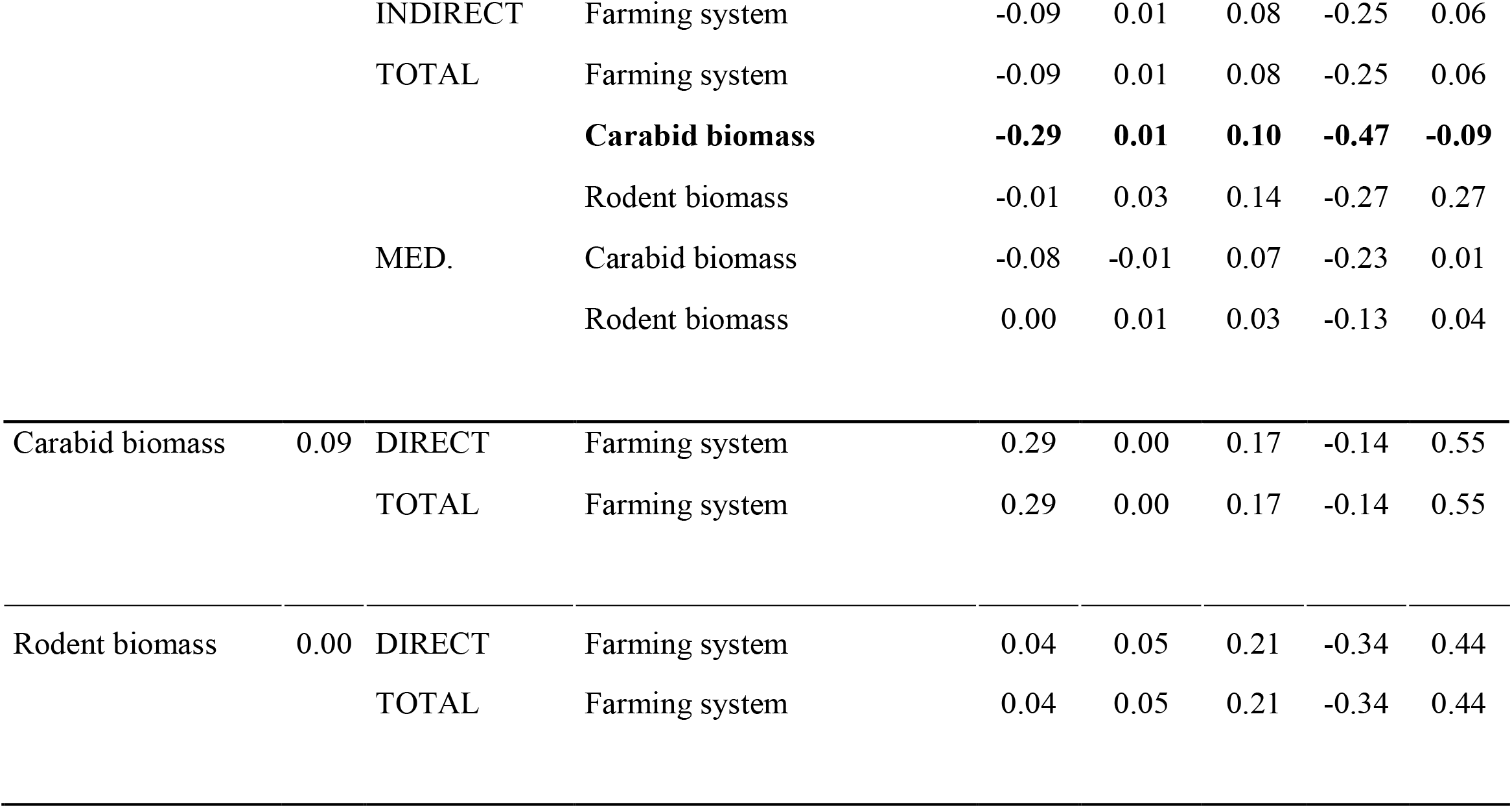
Direct and indirect effects of the predictors based on the retained version of the SEM based on cages with Total Predation. They have been calculated using the semEff package (Murphy, 2022). Five models remain in the final version of the piecewise SEM. Est. means Standardised Estimate, SE means Standard Error, LCI means Lower Confidence Interval, UCI means Upper Confidence Interval and MED. means Mediators. Characters in bold means significative effects.

**Supplementary table 4.** Direct and indirect effects of the predictors based on the retained version of the SEM based on data from cages with Partial Predation. They have been calculated using the semEff package (Murphy, 2022). Four models remain in the final version of the piecewise SEM. Est. means Estimate, SE means Standard Error, LCI means Lower Confidence Interval, UCI means Upper Confidence Interval and MED. means Mediators. Characters in bold means significative effects.

| Response | R<sup>2</sup> | Path | Predictors | Est. | Bias | SE | LCI | UCI |
| --- | --- | --- | --- | --- | --- | --- | --- | --- |
| Yield gap | 0.23 | DIRECT | **Weed biomass** | **-0.48** | **0.01** | **0.17** | **-0.76** | **-0.09** |
|  |  | INDIRECT | **Farming system** | **0.22** | **-0.01** | **0.11** | **0.05** | **0.50** |
|  |  |  | Max n. emerged weeds | 0.06 | 0.00 | 0.09 | -0.07 | 0.28 |
|  |  |  | Carabid biomass | -0.03 | 0.00 | 0.05 | -0.17 | 0.04 |
|  |  | TOTAL | **Farming system** | **0.22** | **-0.01** | **0.11** | **0.05** | **0.50** |
|  |  |  | **Weed biomass** | **-0.48** | **0.01** | **0.17** | **-0.76** | **-0.09** |
|  |  |  | Max n. emerged weeds | 0.06 | 0.00 | 0.09 | -0.07 | 0.28 |
|  |  |  | Carabid biomass | -0.03 | 0.00 | 0.05 | -0.17 | 0.04 |
|  |  | MED. | **Weed biomass** | **0.25** | **0.00** | **0.14** | **0.02** | **0.59** |
|  |  |  | Max n. emerged weeds | -0.04 | 0.00 | 0.06 | -0.25 | 0.04 |
|  |  |  | Carabid biomass | -0.01 | 0.00 | 0.02 | -0.09 | 0.01 |
| Weed biomass | 0.22 | DIRECT | **Farming system** | **-0.47** | **0.03** | **0.16** | **-0.75** | **-0.11** |
|  |  |  | Max n. emerged weeds | -0.12 | 0.02 | 0.17 | -0.42 | 0.24 |
|  |  | INDIRECT | Farming system | 0.02 | -0.01 | 0.04 | -0.02 | 0.12 |
|  |  |  | Carabid biomass | 0.06 | -0.02 | 0.09 | -0.11 | 0.23 |
|  |  | TOTAL | **Farming system** | **-0.45** | **0.02** | **0.15** | **-0.72** | **-0.13** |
|  |  |  | Max n. emerged weeds | -0.12 | 0.02 | 0.17 | -0.42 | 0.24 |
|  |  |  | Carabid biomass | 0.06 | -0.02 | 0.09 | -0.11 | 0.23 |
|  |  | MED. | Max n. emerged weeds | 0.08 | -0.02 | 0.12 | -0.14 | 0.34 |
|  |  |  | Carabid biomass | 0.02 | -0.01 | 0.04 | -0.02 | 0.12 |
| Max n. emerged weeds | 0.27 | DIRECT | **Carabid biomass** | **-0.52** | **0.01** | **0.14** | **-0.74** | **-0.23** |
|  |  | INDIRECT | Farming system | -0.15 | -0.02 | 0.11 | -0.39 | 0.03 |
|  |  | TOTAL | Farming system | -0.15 | -0.02 | 0.11 | -0.39 | 0.03 |
|  |  |  | **Carabid biomass** | **-0.52** | **0.01** | **0.14** | **-0.74** | **-0.23** |
|  |  | MED. | Carabid biomass | -0.15 | -0.02 | 0.11 | -0.39 | 0.03 |
| Carabid biomass | 0.09 | DIRECT | Farming system | 0.29 | 0.01 | 0.17 | -0.19 | 0.55 |
|  |  | TOTAL | Farming system | 0.29 | 0.01 | 0.17 | -0.19 | 0.55 |

**Supplementary Table 5.**
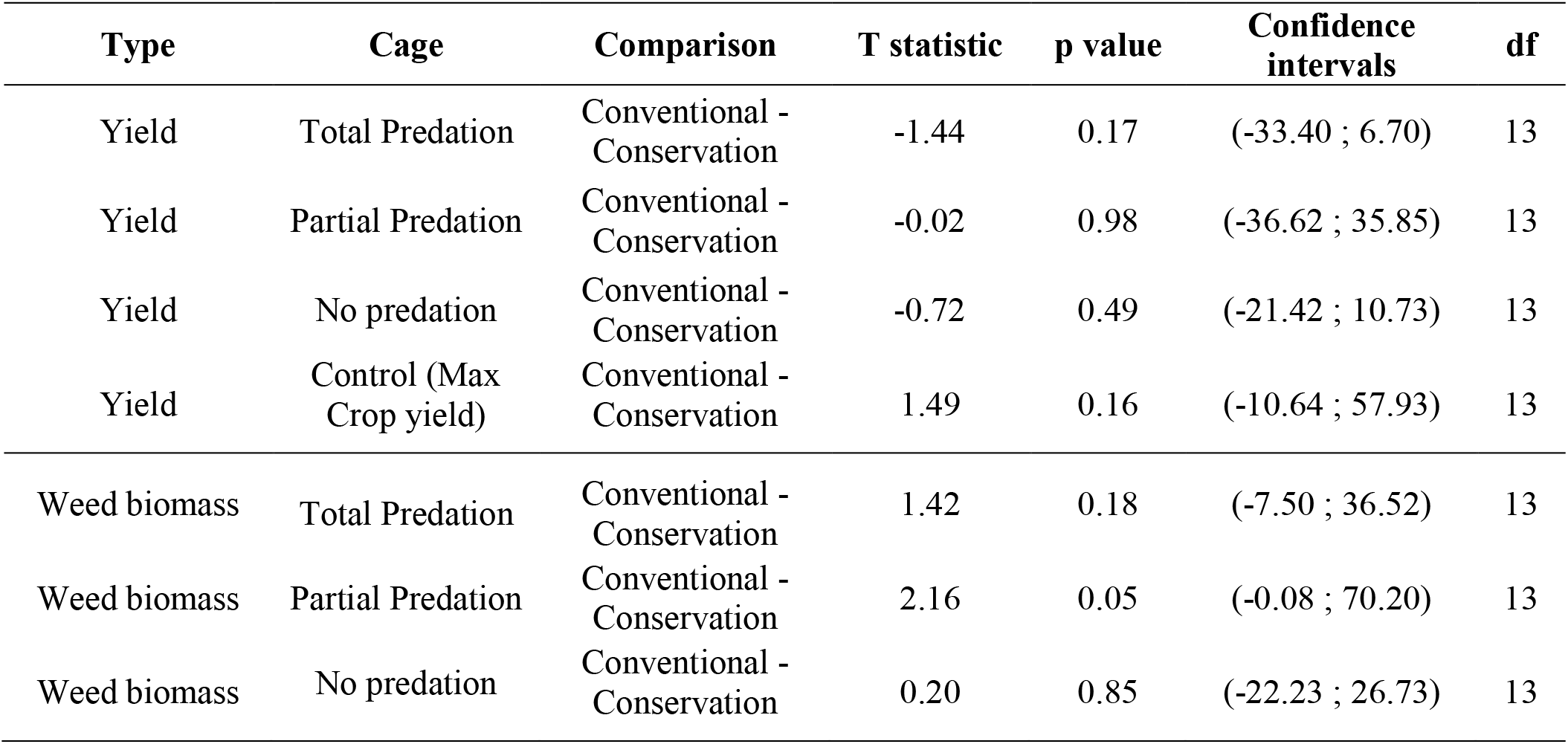
Paired t-tests of comparison of crop yield and weed biomass between conventional and conservation agriculture for each type of cage.

| Type | Cage | Comparison | T statistic | p value | Confidence intervals | df |
| --- | --- | --- | --- | --- | --- | --- |
| Yield | Total Predation | Conventional - Conservation | -1.44 | 0.17 | (-33.40 ; 6.70) | 13 |
| Yield | Partial Predation | Conventional - Conservation | -0.02 | 0.98 | (-36.62 ; 35.85) | 13 |
| Yield | No predation | Conventional - Conservation | -0.72 | 0.49 | (-21.42 ; 10.73) | 13 |
| Yield | Control (Max Crop yield) | Conventional - Conservation | 1.49 | 0.16 | (-10.64 ; 57.93) | 13 |
| Weed biomass | Total Predation | Conventional - Conservation | 1.42 | 0.18 | (-7.50 ; 36.52) | 13 |
| Weed biomass | Partial Predation | Conventional - Conservation | 2.16 | 0.05 | (-0.08 ; 70.20) | 13 |
| Weed biomass | No predation | Conventional - Conservation | 0.20 | 0.85 | (-22.23 ; 26.73) | 13 |

**Supplementary Table 6.** Sampling, seedling, counting and harvesting dates. Counting number of wheat plants and number of tillers per plant in the cages was performed nine times between the dates.

|  |  | Start | End |
| --- | --- | --- | --- |
| Carabid sampling | Sampling date 1 | May-3 2021 | May-5 2021 |
|  | Sampling date 2 | June-2 2021 | June-4 2021 |
| Rodent sampling | Sampling date 1 | May-26 2021 | June-8 2021 |
| Wheat seedling | Seedling date 1 | November-24 2021 | November-29 2021 |
| Weed seed raining mimic | Date 1 | May-17/19 2021 |  |
|  | Date 2 | June-14/16 2021 |  |
| Counting number of wheat plants and number of tillers per plant in the cages | Counting date 1 | June-7 2021 | April-28 2022 |
| Harvesting wheat and weeds in cages and counting plants and tillers | Harvesting date 1 | June-17 2022 | June-30 2022 |

**Supplementary Figure 1.**
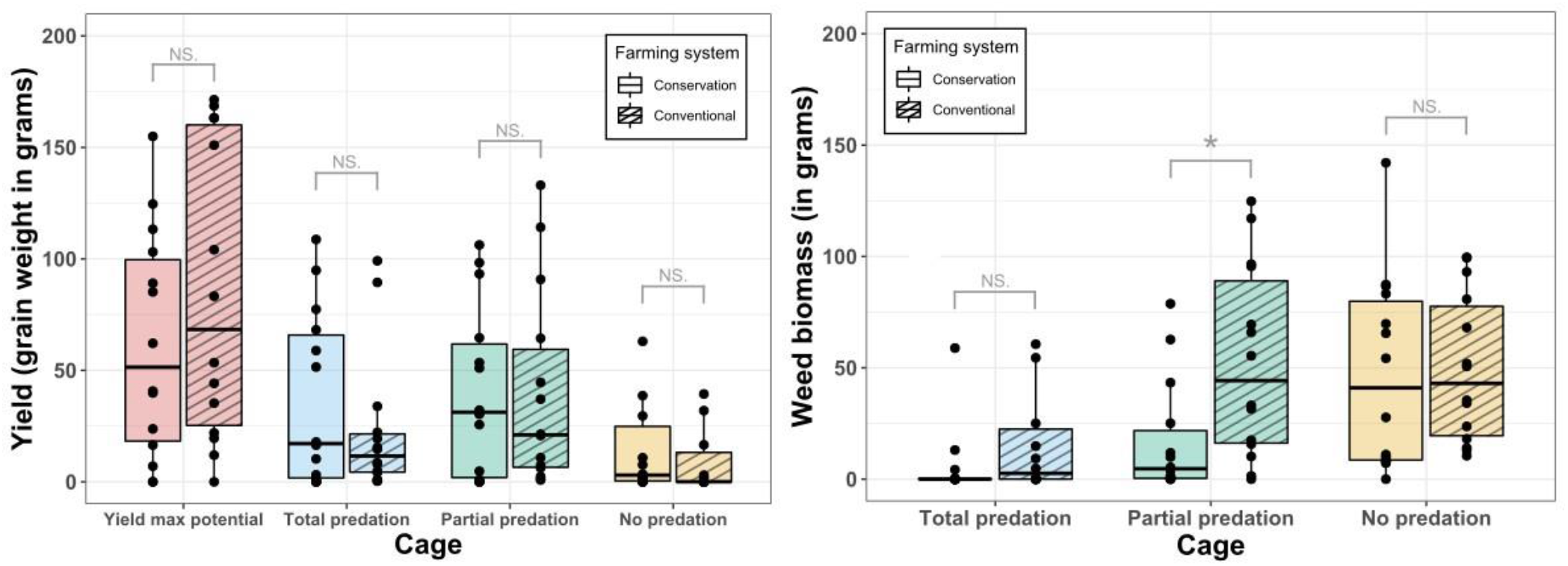
Comparison of crop yield and weed biomass between conventional and conservation agriculture for each cage in the 28 fields. The significance of each comparison is based on paired t-tests (Supplementary table 5).

